# Expanding antimicrobial chemical space by engineering drug safety

**DOI:** 10.1101/2025.10.31.685078

**Authors:** Tahoura Samad, Chayanon Ngambenjawong, Henry Ko, Savan Patel, Cristiana DeAgazio, Heather E. Fleming, Sangeeta N. Bhatia

## Abstract

Antibiotic-resistant (AMR) bacterial infections are a major global health threat. Despite the critical need for new antimicrobials, progress is constrained by protracted development timelines, as well as the requirement for chemical novelty to avoid cross-resistance. Although advances in high-throughput screening, genome mining, and machine learning have greatly accelerated antimicrobial discovery, insufficient separation between antibacterial efficacy and host toxicity remains a bottleneck, precluding the clinical development of many promising compounds. Here, we establish a generalizable, two-component strategy to engineer antimicrobial safety and mobilize otherwise inaccessible chemical space for antimicrobial therapy, using calicheamicin, a potent cytotoxin with unacceptable host toxicity, as a proof of concept. In the first arm, we engineer a conditionally-active drug conjugate that limits calicheamicin activity to infected tissue, thereby reducing systemic toxicity. In the second arm, we co-administer a re-engineered self-resistance enzyme from *Micromonospora echinospora*, the natural producer of calicheamicin, as an “antidote” to neutralize calicheamicin present outside of infected tissue, further mitigating off-target toxicity. The conditionally-active conjugate exhibits activity against Gram-negative and Gram-positive pathogens in response to a protease present within the infected microenvironment. When delivered in combination with the antidote, antibacterial efficacy is maintained while off-target toxicity is reduced in mouse models of Gram positive and negative bacterial pneumonia. We anticipate that our dual strategy, which engineers, rather than selects for enhanced drug safety, by combining conditional drug activity with antidote-driven neutralization of off-target effects, provides a generalizable framework for mobilizing other promising but toxic compounds as antimicrobials.

## INTRODUCTION

Antimicrobial-resistant (AMR) infections are a critical health threat. The World Health Organization (WHO) estimates that resistance to existing drugs directly caused 1.27 million deaths globally and contributed to an additional 4.95 million deaths in 2019 (*1*). Development of new agents to address AMR infections is slow, hindered by the lengthy and costly processes of drug discovery and regulatory approval (*2*), and further challenged by cross-resistance, in which newly developed compounds that are similar to existing antibiotics are subject to the rapid development of resistance (*3*). Additionally, the economics of antibiotic treatment, which can involve short courses of treatment and stewardship programs to limit use, disincentivize investment in antimicrobial development by industry (*2*). Together, these scientific, regulatory, and economic hurdles have resulted in a widening gap between the rise of AMR and the pace of antibiotic development. Given this landscape, there is a critical need for innovative approaches to create antibiotics rapidly, while introducing compounds with novel chemical structures or mechanisms of action that help insulate against cross-resistance.

Recent advances in antibiotic discovery have dramatically expanded our ability to identify compounds with antimicrobial activity. High-throughput screening of small molecule and natural product libraries, machine learning-guided discovery and generation (4-7), large-scale chemical-genetic profiling (8), computational modeling of compound-target interactions (9), and genome mining approaches (10, 11) have increased both the speed and scale of discovery. However, despite this growing discovery capacity, there remains a substantial gap between the number of compounds discovered and those that progress toward clinical development, with advancing candidates often occupying a relatively constrained region of chemical space. A significant reason for this attrition is toxicity; many promising compounds are triaged or exit development early due to a narrow therapeutic index - the difference between the effective therapeutic dose and the dose that causes substantial damage to healthy tissues. Severe toxicities that may be tolerated in other disease indications, such as myelosuppression (16), neuropathy (17), or secondary malignancies (18) are unacceptable for patients with acute infections, who are frequently otherwise healthy. As a result, large portions of the chemical landscape remain underutilized for antibiotic use. Strategies that directly mitigate toxicity therefore represent a powerful opportunity to expand the pool of chemistries accessible for antimicrobial development, enabling the rapid generation of new, structurally diverse antibiotics that carry reduced susceptibility to cross-resistance.

In this study, we present a two-armed, generalizable strategy to widen the therapeutic index of toxic compounds and enable their use as antibiotics. As a proof-of-concept, we apply this strategy to calicheamicin, a potent DNA-damaging agent that is approved for cancer treatment, but which causes significant hepatotoxicity that would preclude administration to patients with infections (*4, 5*). In the first arm of our approach, we formulate calicheamicin as a conditionally-active conjugate, inspired by design principles that have been used to deliver potent cytotoxic payloads in oncology (17,18). In this format, we tether calicheamicin to a carrier protein through a protease-cleavable linker. While conjugated, the cytotoxic activity of calicheamicin is masked until an infection-associated protease releases the drug, thereby confining activity to infected tissue and reducing toxicity to healthy cells, widening the therapeutic index (Fig. 1B). Recognizing that even limited exposure of uninfected tissue to a compound as potent as calicheamicin can cause severe damage, we introduced a second arm into our strategy to neutralize any released, unconjugated drug that travels beyond the infection site. To this end, we adapted a natural microbial strategy by harnessing the self-resistance mechanism of *Micromonospora echinospora*, the natural producer of calicheamicin. *M. echinospora* produces several enzymes that neutralize calicheamicin to protect itself from autotoxicity (*6, 7*); we hypothesized that we could re-engineer and administer these enzymes as “antidotes” to neutralize off-target calicheamicin, further widening the therapeutic index (Fig. 1C). We first show that our calicheamicin-containing drug conjugate exhibits activity dependent on an infection-associated protease against Gram-negative and -positive priority pathogens, and that prior to cleavage, this conjugated format masks calicheamicin toxicity to mammalian cells. We further show that bacterial-derived calicheamicin self-resistance enzymes can be re-engineered to act as long-circulating antidotes that neutralize off-target calicheamicin *in vitro* and *in vivo*, reducing liver toxicity. Finally, we validated that the combination of these two strategies achieves therapeutic efficacy in mouse models of Gram positive and negative bacterial pneumonia, while minimizing systemic toxicity. Together, these results establish a generalizable framework for engineering drug safety and mobilizing potent but toxicity-limited compounds as antimicrobials.

**Figure 1.**
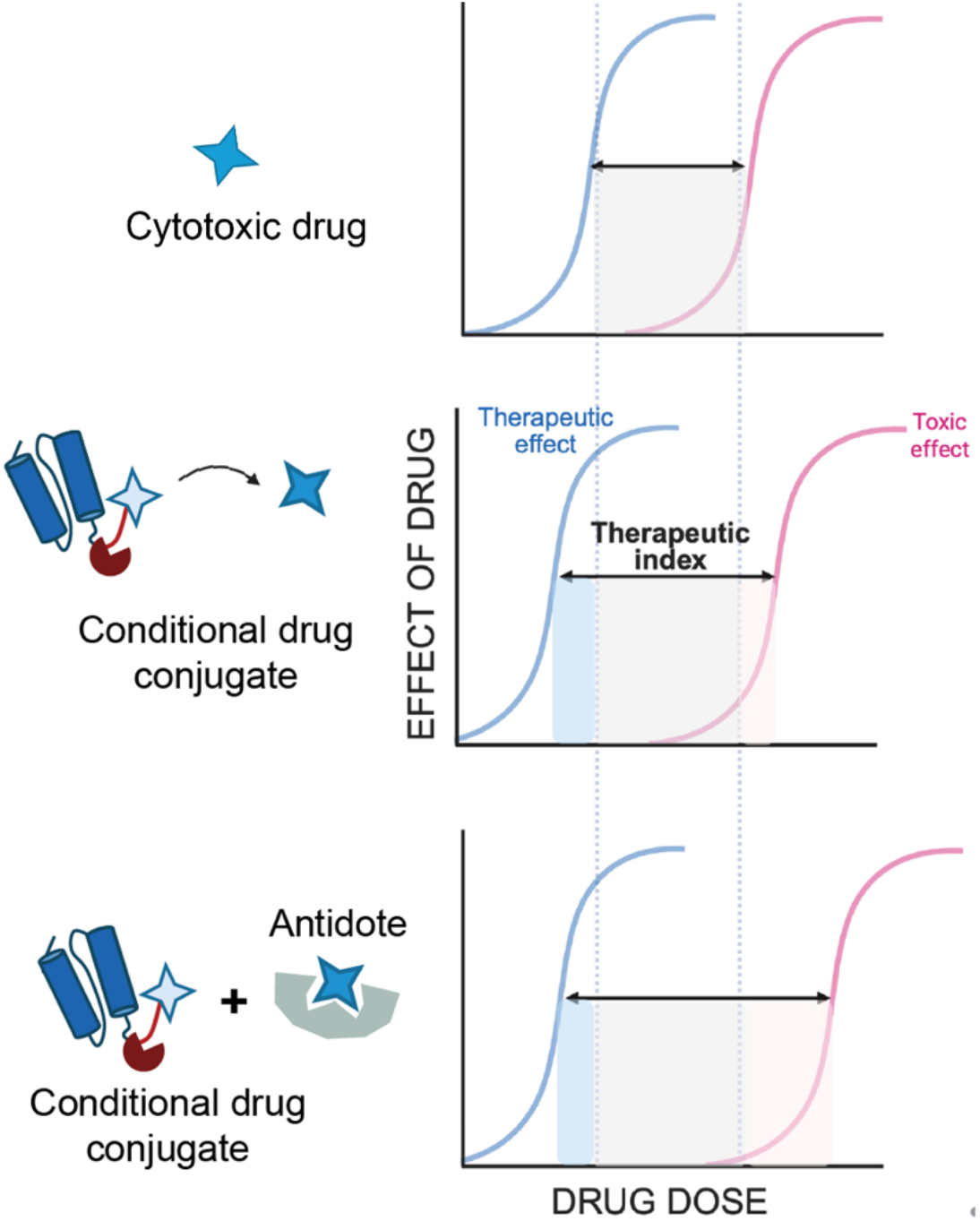
Dual strategy to expand the therapeutic index of toxic compounds for antimicrobial use. (Top) A narrow therapeutic index, defined as the difference between the median therapeutic effect and the median toxic effect, constrains the utility of cytotoxins (blue star) as antimicrobials. (Middle) Formulation of cytotoxins as conditionally active conjugates can widen the therapeutic index by restricting drug activity to an infection site, where proteases (red pac-man) release drug from the conjugate. (Bottom) “Antidotes” derived from self-resistance enzymes further widen the therapeutic index by neutralization off-target drug.

## RESULTS

### Calicheamicin has broad spectrum activity against Gram-positive and Gram-negative pathogens

We first evaluated the antimicrobial activity of calicheamicin against a panel of Gram-negative and Gram-positive bacterial pathogens, including several designated as priority pathogens by the World Health Organization (*8*). Measuring microbial susceptibility to calicheamicin using broth microdilution assays, we determined the minimum inhibitory concentration (MIC) of calicheamicin across representative strains (Table 1). We found that calicheamicin had potent activity against the Gram-positive bacterium *Staphylococcus aureus*, with an MIC of 0.0006 µg/mL, a potency that was comparable to or greater than that of a clinically used antibiotic vancomycin (Table 1). Notably, this potency was maintained against a methicillin-resistant *S. aureus* (MRSA) isolate, BA1556. Calicheamicin also demonstrated activity against Gram-negative pathogens including *Pseudomonas aeruginosa, Escherichia coli*, and *Klebsiella pneumoniae*, with MIC values ranging from 0.015 to 0.5 µg/mL (Table 1). Together, these data demonstrate that calicheamicin exhibits broad-spectrum antimicrobial activity, with efficacy extending to multidrug-resistant strains, and at concentrations that would be realistic for clinical translation.

**Table 1.**
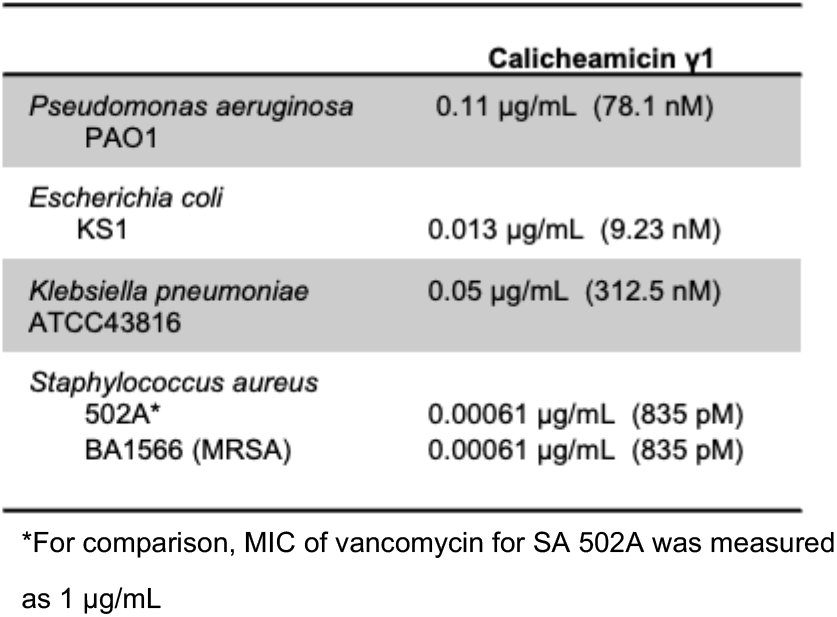
Antibacterial activity of calicheamicin against representative Gram-negative and Gram-positive pathogens. Minimum inhibitory concentrations (MICs) were determined by microdilution assay in Mueller– Hinton medium supplemented with 3% DMSO. Data represent biological triplicates.

### Conditionally-active drug conjugate masks cytotoxicity while retaining antibacterial activity *in vitro*

While the potent antimicrobial activity of calicheamicin is a promising starting point, its substantial toxicity toward mammalian cells, particularly liver cells, precludes its use as a free compound. In oncology, this limitation has been addressed by formulating calicheamicin as an antibody–drug conjugate (*9, 10*), where targeting and conditional activity improve the therapeutic index. Building on this approach, we developed a conditionally-active calicheamicin conjugate tailored for the infection context. In this design, calicheamicin was tethered to an albumin-binding domain (ABD) via a protease-sensitive linker that masks its cytotoxic activity until cleavage by infection-associated proteases (Fig. 2A). Use of albumin as the carrier protein leverages endogenous trafficking pathways to enhance accumulation (*11, 12*), retention (*13*), and mucopenetration (*14*) at sites of infection. As a linker between albumin and the drug, we used a peptide sequence cleavable by neutrophil elastase, which is enriched at sites of bacterial infection due to immune cell recruitment.

**Figure 2.**
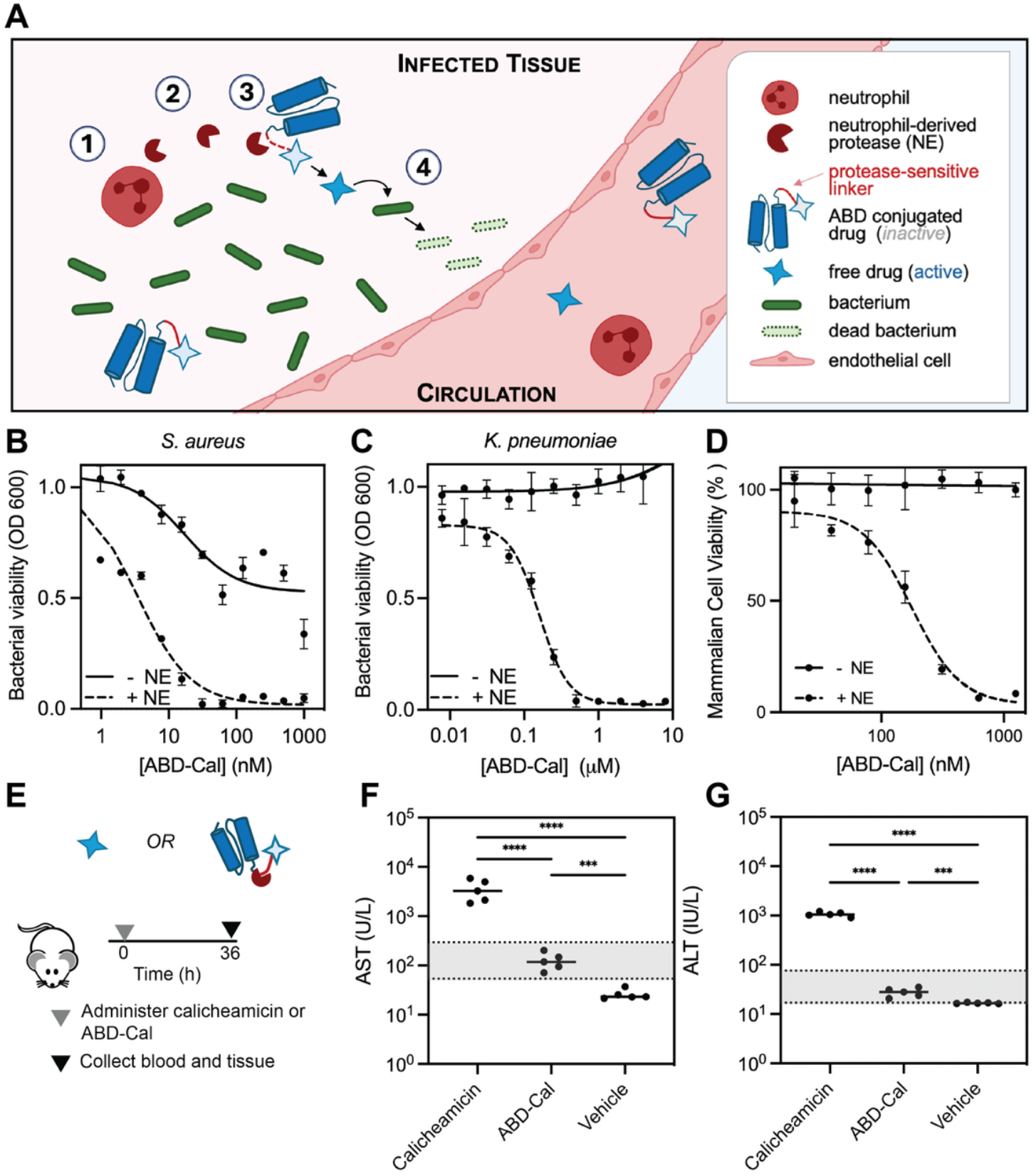
ABD-calicheamicin conjugate demonstrates masked mammalian toxicity and protease-triggered antibacterial activity *in vitro* and improved safety *in vivo*. (A) Schematic of ABD-calicheamicin conjugate (ABD-Cal) activation within the infection microenvironment. Within the infection environment, 1) neutrophils are recruited and 2) secrete neutrophil elastase which 3) cleaves and releases calicheamicin from the conjugate 4) such that it can kill bacteria. (B-C) Antimicrobial activity of ABD-Cal evaluated via microdilution assays against representative (B) Gram-positive and (C) Gram-negative bacterial strains in the presence or absence of recombinant neutrophil elastase. Bacterial growth measured by absorbance at 600 nm and normalized to untreated controls. Fold change in antimicrobial activity calculated as the ratio of MIC values of intact vs. cleaved conjugates. (mean ± sd, n = 3 biological replicates) (D) Masked mammalian toxicity of ABD-Cal evaluated via MTS cell proliferation assay in HepG2 cells. Cell viability measured by absorbance at 490 nm and normalized to untreated controls. Fold change in cytotoxicity calculated as the ratio of IC_50_values of intact vs. cleaved conjugates. (mean ± sd, n = 3 biological replicates) (E) Schematic of experimental timeline of mice administered free calicheamicin (0.025 mg/kg) compared with ABD-Cal (0.025 mg/kg drug eq.) (F) Serum AST and (G) ALT. Dashed lines indicate normal healthy range. (n = 5 biological replicates. Statistical significance was determined by one-way ANOVA followed by Tukey’s post hoc test). *P < 0*.*001 (\*\*\**); *P* < 0.0001 (****)

To generate the ABD–calicheamicin conjugate (ABD-Cal), we prepared a calicheamicin–peptide intermediate and coupled it to a recombinantly-expressed ABD using strain-promoted azide–alkyne cycloaddition. We next tested whether conjugation masked the cytotoxic activity of calicheamicin and allowed for protease-dependent activation by assessing antibacterial activity and mammalian cell toxicity in the presence or absence of neutrophil elastase *in vitro*. The conjugate was incubated with neutrophil elastase or vehicle control for 4 hours prior to evaluation. We then measured antibacterial activity against a representative Gram-positive (*S. aureus*) (Fig. 2B) and Gram-negative (*K. pneumoniae*) pathogen (Fig. 2C). In the absence of protease, we observed no antimicrobial activity; in the presence of protease, we observed a recovery in antimicrobial activity with a (∼60-120-fold) change in the minimum inhibitory concentration (MIC) between the intact and cleaved conjugates, depending on the pathogen tested. We also assessed whether formulation of calicheamicin as a conditionally-active conjugate protected mammalian cells from toxicity. We found that in the absence of protease, the intact conjugate demonstrated minimal mammalian toxicity to HepG2 liver cells after 24 hours of exposure (Fig. 2D). Together, these data show that calicheamicin, when formulated as a conditional conjugate, exhibits reduced toxicity toward mammalian cells while conjugated, but regains potent antibacterial activity once released by protease cleavage.

### Conditionally-active drug conjugate demonstrates improved safety relative to free drug *in vivo*

Having shown *in vitro* that conjugation masks calicheamicin’s cytotoxicity and that protease cleavage restores antibacterial activity, we next asked whether restricting activation to the infection microenvironment reduces systemic, particularly hepatic, toxicity *in vivo*. To test this, we compared the liver toxicity of the conditionally active calicheamicin conjugate with free calicheamicin *in vivo*. Healthy mice were intravenously administered either free calicheamicin or the conjugate (dose matched by drug concentration) (Fig. 2E). Liver injury was assessed 36 h post-injection by measuring serum aspartate aminotransferase (AST) levels (Fig. 2F), alanine aminotransferase (ALT) (Fig. 2G) and by histological analysis (Fig. S1). Mice treated with free calicheamicin exhibited elevated AST (Fig. 2F) and ALT (Fig. 2G) levels. Mice receiving the conjugate showed a significant reduction in toxicity relative to free drug, although serum liver enzymes were significantly elevated above the baseline of mice treated with vehicle alone. Histology revealed moderate to severe injury in mice treated with free drug, evidenced by diffuse presence of hepatocytes with swollen cytoplasm or shrunken, hypereosinophilic cells (Fig. S1) The livers of conjugate-treated mice showed more mild changes, including some apoptotic cells with swollen cytoplasm, and some healthy cells, consistent with more limited toxicity (Fig. S1). These findings indicate that formulation of calicheamicin as the conditionally-active conjugate significantly reduces off-target hepatotoxicity *in vivo*.

### Microbial-derived antidotes can neutralize drug toxicity *in vitro* and *in vivo*

Formulation of calicheamicin as a conditional conjugate significantly reduced hepatotoxicity in healthy mice but did not eliminate liver injury, which we hypothesized arose from a fraction of drug released outside the infection site, potentially via low levels of protease-mediated activation outside the infection environment, or diffusion of released drug from infected tissue back into the bloodstream. We therefore developed a second complementary strategy to address these residual sources of toxicity and further improve the safety profile of calicheamicin. To achieve a second layer of toxicity mitigation, we drew inspiration from the natural self-resistance mechanism of *Micromonospora echinospora*, the native producer of calicheamicin, which avoids self-toxicity by enzymatically inactivating the compound. We envisioned adapting this self-resistance strategy for therapeutic use by engineering long-circulating variants of these calicheamicin-neutralizing enzymes to act as “antidotes”, intercepting and neutralizing calicheamicin that escapes the infection microenvironment before it reaches sensitive tissues (Fig. 3A).

**Figure 3.**
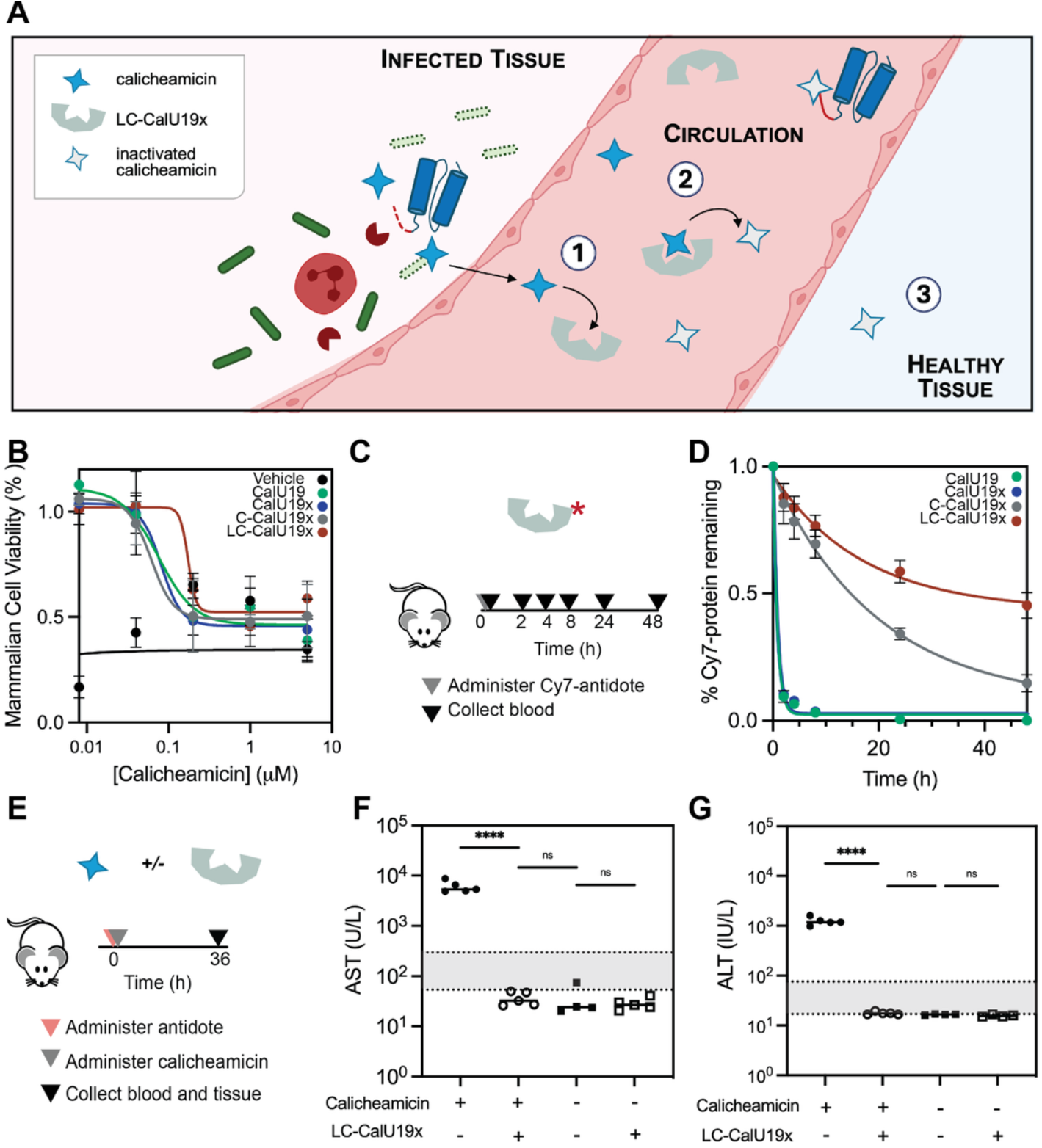
Microbial-inspired antidotes protect against calicheamicin-mediated liver toxicity *in vitro* and *in vivo*. Schematic illustrating neutralization of calicheamicin by engineered microbial self-resistance enzymes (“antidotes”). After neutrophil-elastase–mediated cleavage of the ABD–calicheamicin conjugate, either on-target (within the infection microenvironment) or off-target (within healthy tissue) released calicheamicin can 1) enter circulation, where it is 2) bound and inactivated by LC-CalU19x, such that exposure of healthy tissue to active drug is reduced. (B) Calicheamicin neutralization by engineered variants of CalU19 (evaluated via cell proliferation assay in HepG2 cells. Cell viability measured by absorbance at 490 nm and normalized to untreated controls. mean ± sd, n = 4 biological replicates. (C) Schematic of timeline during which Cy7-conjugated proteins were assayed for clearance from plasma. (D) Pharmacokinetics of antidote variants following systemic injection, with plasma concentrations of Cy7 quantified over time. (mean ± sd, n = 4–5 mice per group) (E) Schematic of *in vivo* neutralization of free calicheamicin by LC-CalU19x.(F) Serum AST and (G) ALT levels measured 36 h after calicheamicin administration with or without antidote treatment. Dashed lines indicate normal healthy range (n = 5 Statistical significance determined by one-way ANOVA followed by Tukey’s post hoc test.) *P* < 0.0001 (****); ns: not significant.

We first evaluated which of the native self-resistance enzymes would be most suitable for adaptation as a circulating antidote. The calicheamicin biosynthetic cluster encodes three self-resistance enzymes, CalC, CalU16, and CalU19 (*6, 7*). Reduction of calicheamicin’s trisulfide bond generates the reactive species responsible for its cytotoxicity. CalC and CalU16 act only on this reduced form, whereas CalU19 also neutralizes the unreduced molecule, making it best suited for calicheamicin neutralization in circulation, which is not a reducing environment. We recombinantly expressed CalU19 and sought to confirm its neutralizing activity *in vitro*, by evaluating if administration of CalU19 could protect mammalian cells from calicheamicin-induced toxicity. HepG2 cells were incubated with serial dilutions of calicheamicin in the presence or absence of a fixed concentration of CalU19 for 24 hours, and cell viability was assessed by MTS assay (Fig. 3B). CalU19 conferred protection, with the highest protection observed at lower calicheamicin concentrations, consistent with the stoichiometric mechanism of inactivation in which one molecule of CalU19 is consumed per molecule of calicheamicin neutralized (Fig. 3B).

Following *in vitro* validation of CalU19 activity, we next assessed its ability to selectively neutralize free calicheamicin without affecting the conjugated drug. During calicheamicin neutralization, CalU19 undergoes a unique self-cleavage event, which can be used as an indicator of neutralizing activity. SDS-PAGE of CalU19 incubated with calicheamicin, ABD-Cal or vehicle, revealed CalU19 self-cleavage fragments only when CalU19 was incubated with free calicheamicin (Fig. S2), confirming that neutralizing activity is restricted to free drug.

To realize our design goal of a long-circulating antidote capable of soaking up off-target drug *in vivo*, we next engineered CalU19x for *in vivo* administration. We first removed two unpaired cysteines, which could promote off-target interactions with other proteins in vivo, substituting them with alanine to generate CalU19x, which retained neutralizing activity comparable to the wild-type enzyme (Fig. 3B). We then assessed CalU19 *pharmacokinetics in vivo* (Fig. 3C, D). Fluorophore-labeled CalU19 and CalU19x were both rapidly cleared from circulation, with plasma half-lives of ∼1 h. To extend the circulation half-life of CalU19x, we generated fusions to single (C-CalU19x) and tandem (LC-CalU19x) albumin-binding domains, as these extend half-life through association with serum albumin. LC-CalU19x exhibited the longest half-life, remaining detectable for at least 48 h in plasma (Fig. 3D), and maintained neutralizing activity *in vitro* comparable to wild-type CalU19 (Fig. 3B).

Having established a long-circulating antidote *in vitro*, we next evaluated whether LC-CalU19x could protect against calicheamicin toxicity *in vivo*. Healthy animals were intravenously administered LC-CalU19x or vehicle followed by dosing with calicheamicin (Fig. 3E). Serum and tissue were collected 36-48 h later, and serum AST and ALT levels were quantified. Mice administered calicheamicin and LC-CalU19x showed significantly reduced serum AST (Fig. 3F) and ALT (Fig. 3G) compared with mice administered calicheamicin and vehicle, indicating mitigation of calicheamicin-induced hepatotoxicity. Histology (Fig. S3) agreed with these findings: livers from mice administered calicheamicin and LC-CalU19x-treated mice were largely indistinguishable from healthy controls, whereas mice treated with calicheamicin alone displayed hepatocellular degeneration and necrosis, indicated by swollen cells with eosinophilic cytoplasm (Fig. S3). Together, these results demonstrate that LC-CalU19x acts as a circulating antidote that mitigates off-target toxicity without interfering with the activity of the conditionally-active calicheamicin conjugate, supporting their use as a combined therapy.

### Drug conjugate and antidote combination achieves antimicrobial efficacy in mouse models of pneumonia

We next evaluated the preclinical potential of the ABD-calicheamicin conjugate and LC-CalU19x in combination, testing whether co-administration of the antidote with the conditionally-active drug could reduce bacterial burden while minimizing liver toxicity in a mouse models of bacterial pneumonia. Mice were infected with *S. aureus*, followed by intratracheal administration of vehicle, ABD-calicheamicin alone, ABD-calicheamicin in combination with intravenously administered LC-CalU19x (Fig. 4A–B), or intravenously administered vancomycin. Treatment of *S. aureus*-infected mice with the conditionally-active conjugate with or without antidote resulted in a 4-log reduction in lung bacterial burden 6 h compared with the vehicle control, demonstrating that on-target antimicrobial efficacy is preserved in the presence of the antidote (Fig. 4B). At the same timepoint, treatment with vancomycin produced a 1–2-log reduction in bacterial burden (Fig. 4B). To assess safety, we compared liver toxicity between mice treated with ABD-Cal alone or ABD-Cal with LC-CalU19x. We observed a significant reduction in toxicity in mice receiving the conjugate plus antidote relative to the conjugate-only and vehicle groups, as measured by serum AST (Fig. 4C) and ALT (Fig. 4D), with enzyme levels in antidote-treated mice within the healthy range.

**Fig. 4.**
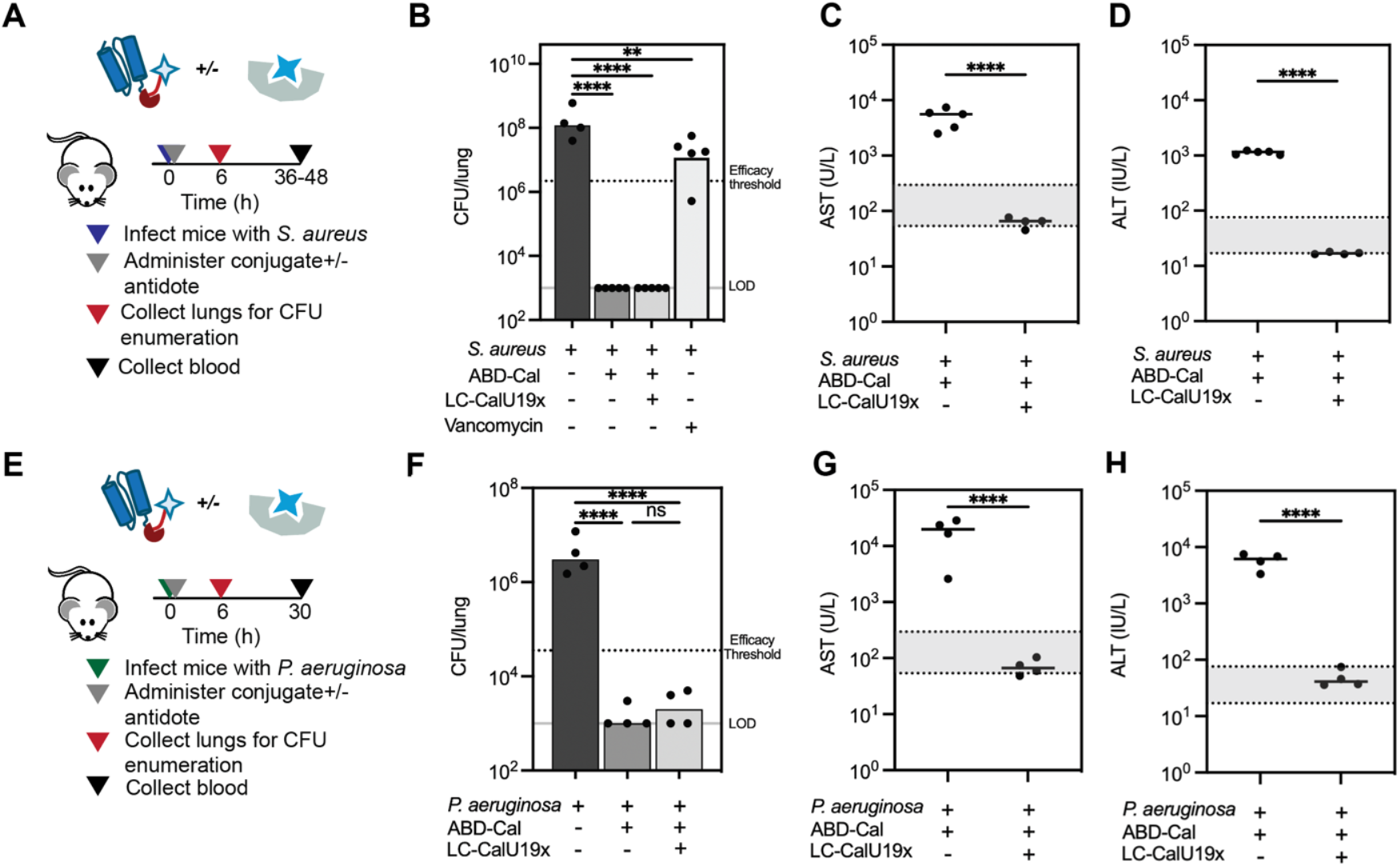
Therapeutic efficacy of calicheamicin conjugate and neutralizing antidote in a murine models of Gram positive and negative bacterial pneumonia. (A) Schematic of experimental timeline for treatment of *S. aureus* pneumonia. For lung bacterial burden and serum chemistry analyses, separate cohorts of animals were used for tissue or blood collection at the indicated time points. (B) Bacterial burden in lungs 6 h post-treatment in mice infected with *Staphylococcus aureus* 502A and treated intratracheally with vehicle, treated intratracheally with calicheamicin drug conjugate (0.25 mg/kg calicheamicin eq) (ABD-Cal) and then intravenously with vehicle or antidote (LC-CalU19x), or treated intravenously with vancomycin (110 mg/kg). Each point represents an individual animal (n = 4-5 per group). Dashed line indicates the threshold for efficacy, defined as a two-log reduction in bacterial burden from the vehicle treated group. Grey line indicates the limit of detection for CFU quantification. Statistical significance determined by one-way ANOVA followed by Dunnett’s post hoc test. (C) Serum levels of AST and (D) ALT measured 36-48 h after infection and treatment with ABD-Cal, in the presence or absence of antidote. Each point represents an individual animal (n = 4–5 per group). Dashed lines indicate the normal healthy range. Statistical significance was determined by unpaired t-test. (E) Schematic of experimental timeline for treatment of *P. aeruginosa* pneumonia. (F) Bacterial burden in lungs 6 h post-treatment in mice infected intratracheally with *Pseudomonas aeruginosa* PAO1 and treated intratracheally with vehicle or ABD-Cal (0.4 mg/kg calicheamicin eq) and then intravenously with vehicle or antidote (LC-CalU19x). Each point represents an individual animal (n = 4 per group). Statistical significance determined by one-way ANOVA followed by Tukey’s post hoc test. (G) Serum levels of AST and (H) ALT measured 30 h after infection and treatment with ABD-Cal, in the presence or absence of antidote. Each point represents an individual animal (n = 4 per group). Dashed lines indicate the normal healthy range. Statistical significance was determined by unpaired t-test. P < 0.01 (**); P < 0.001 (***); P < 0.0001 (****).

To evaluate the generalizability of this approach across pathogens, we additionally tested the drug conjugate and antidote combination in a mouse model of Gram-negative *Pseudomonas aeruginosa* pneumonia (Fig. 4E). In this model, treatment with the drug conjugate with and without antidote resulted in a 3-log reduction in lung bacterial burden relative to vehicle control at 6 h (Fig. 4F). Consistent with the findings in the *S. aureus* model, co-administration of the antidote significantly reduced liver toxicity compared with treatment with conjugate alone, as measured by serum AST and ALT (Fig. 4G–H). Together, these findings demonstrate that pairing a conditionally-active drug conjugate with a long-circulating antidote preserves potent antimicrobial efficacy while substantially reducing systemic toxicity. Furthermore, the conditional drug–antidote strategy is effective across both Gram-positive and Gram-negative pathogens, supporting the potential broad applicability of this platform for the treatment of bacterial infections.

## DISCUSSION

In this work, we develop a generalizable, two-component strategy designed to widen the therapeutic index of potential antimicrobial compounds whose clinical translation is otherwise limited by toxicity. Using calicheamicin, a cytotoxin with antimicrobial activity but unacceptable hepatotoxicity, as a proof of concept, we demonstrate that antibacterial activity and host toxicity can be decoupled through the combined use of conditional drug activation to restrict drug activity to infected tissue and active detoxification of off-target drug. Together, this approach enabled therapeutic efficacy in a mouse model of bacterial pneumonia while minimizing hepatotoxicity.

Conditional drug conjugates hold considerable potential for infectious disease, but their development in that context is far less common than in oncology, where antibody–drug conjugates (ADCs) have achieved notable clinical success (*11, 12, 15, 16*). The distinct biological context of bacterial infections requires revising design criteria that have guided drug conjugate development in oncology. While oncology relies on antibodies to direct cytotoxic payloads to defined tumor antigens, infection-focused conjugates would benefit from carrier proteins that target the site of infection in an antigen-independent manner and are thus compatible with broad-spectrum activity against a variety of pathogens. Our design (Fig. 2) uses an albumin-binding domain (ABD) as the drug carrier, with the intent to promote retention (*13*) and mucopenetration (*14*) at the sites of infection when administered into the lung. For the linker connecting the carrier protein to the drug, design requirements differ substantially between cancer and bacterial infection. ADCs for cancer typically exploit tumor-associated proteases such as cathepsins to release payloads intracellularly within lysosomes, a defined environment with acidic pH, reducing conditions, and a well-characterized protease repertoire (*17*). In contrast, bacterial infection is likely to require extracellular drug release, which is a heterogeneous environment, where disease-associated triggers act amid variable pH and redox states in the presence of immune cells and cellular debris. To achieve robust and selective activation in such a setting, we incorporated a peptide linker cleavable by neutrophil elastase, which is enriched extracellularly at sites of bacterial infection after release by recruited immune cells (*13*).

Formulation of calicheamicin as a conditionally-active conjugate reduced liver toxicity but did not fully eliminate it (Fig. 2), consistent with both preclinical and clinical data from oncology, the context in which conditional-drug conjugates have advanced the furthest clinically. In particular, published evidence suggests that the maximum tolerated doses of some ADCs approach those of the unconjugated payload (*5, 18*–*20*). Prior efforts have focused on refining aspects of the antibody, linker, or payload to reduce toxicity, but off-target damage in the liver, bone marrow, and gastrointestinal tract remains a major barrier. To move beyond these limitations in the context of infection, we turned to microbes themselves, which not only produce some of the natural products we used as therapeutics, but also encode self-resistance enzymes to protect against autotoxicity. Inspired by this microbial strategy, we introduced a second layer of protection: delivery of an engineered self-resistance enzyme derived from *Micromonospora echinospora*, the natural producer of calicheamicin, as an antidote to neutralize circulating free drug outside the site of infection (Fig. 3). While coadministration of antibiotics with compounds that enhance antimicrobial activity are under development (*21*–*24*), coadministration of agents specifically designed to improve therapeutic safety is less explored in infectious disease. Related coadministration strategies aimed at improving therapeutic safety have begun to be explored in oncology contexts, for example in preclinical work examining neutralization of toxicity for radiation (*25*), and clinically in the administration of agents (*26*) to mitigate cytokine release syndrome after administration of CAR-T therapy.

Despite the promise of our drug conjugate and antidote platform, several challenges remain before its translation to clinical settings. Microbial-derived proteins like the antidote we developed may be immunogenic upon repeated administration, as is often required in the treatment of bacterial infections. Existing molecular engineering strategies such as PEGylation (*27*), domain humanization (*28*), or epitope masking (*29*) of the antidote could reduce its immunogenic potential, and there are encouraging examples of regulator-approved and clinically-used products derived from microbial enzymes (*30, 31*). Second, the success of this dual system hinges on spatial, kinetic, or temporal separation between on-target activity and off-target detoxification. In our current model, we achieve this distinction using physical compartmentalization, via local pulmonary delivery of the drug conjugate and systemic administration of the antidote (Fig 4). This separation between localized activity and off-target neutralization is also achieved by virtue of the conjugate, which protects the conjugated drug from premature deactivation by the antidote, as the enzyme can only bind and neutralize the released, active form of the drug (Fig. S2). However, extending the paired conjugate-antidote system to other sites of infection will require context-specific tailoring. For example, we hypothesize that treatment of systemic bloodstream infections would likely require prophylactic delivery of CalU19, either as a protein or mRNA encapsulated in lipid nanoparticles, such that they can be taken up by liver sinusoidal endothelial cells or hepatocytes to protect these tissues prior to systemic administration and release of calicheamicin from the conjugate in response to cues from the bloodstream infection. The modularity of our platform, with the ability to exchange the carrier protein and protease-susceptible linker components of the conjugate, as well as the possibility of fusing other targeting partners to CalU19 to control half-life or localization within the body, makes such adaptation feasible. Finally, the kinetics of enzyme-mediated neutralization of “off target” drug relative to bacterial uptake and killing by “on target” drug remain incompletely understood. Future studies may benefit from structure-guided engineering or directed evolution of CalU19 to modulate its catalytic efficiency to further enhance the therapeutic window of the conjugate and antidote pair.

Beyond calicheamicin, we propose that this dual strategy—(1) formulating drugs as conjugates to achieve selective activity at the site of infection and (2) pairing them with microbial resistance enzymes that neutralize off-target toxicity—represents a generalizable approach for mobilizing antimicrobial compounds whose development or use is limited by toxicity, such as existing antibiotics with narrow therapeutic windows, investigational compounds previously abandoned because of safety or pharmacokinetic liabilities, and newly discovered molecules emerging from genome-mining or machine-learning–driven discovery pipelines(*32*–*35*). Drug conjugates are modular: infection-targeting carrier proteins, conditional linkers, and payloads can be interchanged to suit different pathogens and infection contexts(*36*). Further, microbial resistance mechanisms provide a large reservoir of potential antidotes. Many natural product biosynthetic gene clusters encode self-resistance genes that protect the producing organism from autotoxicity(*37*); related resistance determinants can also be found in organisms within the same ecological communities as natural product producers (*38*) . These genes encode enzymes that can chemically modify, degrade, or sequester a compound, detoxification mechanisms that could be repurposed therapeutically. Advances in genome mining and machine learning could accelerate the discovery and prioritization of additional drug–antidote pairs. Beyond mining natural mechanisms, these same tools may also enable the de novo design of antidotes (*39, 40*), providing additional routes to engineer safety for antimicrobial compounds.

.More broadly, the combined use of the drug conjugate and antidote represents a conceptual shift in antibiotic treatment—towards a form of molecular stewardship—in which each molecule of the antimicrobial is accounted for, with on-target drug directed precisely to the site of infection while off-target drug is actively neutralized. Although mitigation of liver toxicity was a primary goal of this work, the same principle could reduce collateral damage to tissues that are commonly sensitive to cytotoxic agents, such as the bone marrow. Antibiotic-induced dysbiosis of the gut microbiome is another major concern; antibiotic treatment is frequently associated with shifts in microbial composition, overgrowth of opportunistic pathogens such as *Clostridioides difficile*, and impaired gut barrier function (*41, 42*). We hypothesize that co-administration of an antidote could attenuate antibiotic-associated microbiome disruption by neutralizing drug before it reaches the gastrointestinal tract. Beyond the host, excreted antibiotics that enter into the environment through sewage and wastewater exert selective pressure on environmental bacteria, which can drive the emergence of resistance through exposure to sublethal concentrations (*43*). The excretion of highly potent cytotoxic compounds such as calicheamicin also raises concerns about ecological toxicity (*44*–*46*). Incorporation of antidotes into therapeutic regimens may therefore not only improve safety within the patient but also mitigate the broader ecological consequences of antimicrobial use. Finally, outside of the context of infectious disease, we envision antidotes as an enabling technology for conditionally-active conjugates more generally, with the potential to mitigate toxicity in oncology and other therapeutic indications where drugs with potentially severe side effects and dose limiting toxicities are administered.

## ACKNOWLEDGEMENTS, FUNDING AND COMPETING INTERESTS

We are grateful to Dr. Laura Kiessling and Dr. Deborah Hung for their scientific insight and support over the course of the work. We are grateful to the Koch Institute Swanson Biotechnology Center, specifically the Histology core (Kathy Cormier, Magalie Boucher) and the Biopolymer and Proteomics core (Heather Amoroso). We are grateful to Kasturi Chakraborthy, Susanna Elledge and Vardhman Kumar for feedback on the manuscript. This work was supported by R01 AI132413 and U19 AI142780 grants from the National Institute of Allergy and Infectious Diseases. This study was supported in part by the Virginia and D.K. Ludwig Fund for Cancer Research, the Koch Institute Support Grant P30-CA14051 from the National Cancer Institute (Swanson Biotechnology Center) and Core Center Grant P30-ES002109 from the National Institute of Environmental Health Sciences. Additional support was received from the Koch Institute’s Marble Center for Cancer Nanomedicine. This research was supported in part by a generous gift from Joel Gantcher. T.S. was supported by a postdoctoral fellowship from the Ludwig Center at MIT’s Koch Institute for Integrative Cancer Research. S.N.B. is a Howard Hughes Institute Investigator. Competing interests: S.N.B. reports consulting roles and/or equity in Amplifyer Bio, Catalio Capital, Danaher, Earli Inc., Impilo Therapeutics, Matrisome Bio, Ochre Bio, Port Therapeutics, Satellite Bio, Sunbird Bio, and Xilio Therapeutics, which were not involved in this study.

## MATERIALS and METHODS

### Bacterial strains and culture

*Escherichia coli* ATCC 29522, *Klebsiella pneumoniae* ATCC 43816, *Pseudomonas aeruginosa* PAO1, *Staphylococcus aureus* 502A, and *Staphylococcus aureus* USA300_FPR3757 were used in this study. Strains were obtained from the American Type Culture Collection (ATCC) or as indicated. Bacteria were cultured in Mueller–Hinton broth (MHB; BD Difco) at 37 °C with shaking at 250 r.p.m. Growth was monitored by optical density at 600 nm (OD600).

### Molecular cloning

gBlock™ gene fragments encoding ABD with a C-terminal cysteine ((ABD)_2_-(EEG)_6_-Cys), CalC, and CalU19 with flanking restriction sites NcoI and XhoI were ordered from Integrated DNA Technologies, cloned into pET28(a) by restriction enzyme digestion and ligation, and transformed into *E. coli* DH5α competent cells (New England Biolabs). Colonies encoding the correct inserts were confirmed by Sanger sequencing (QuintaraBio), and plasmids were purified using a plasmid miniprep kit (Thermo Fisher Scientific). For CalU19 variants, clonal genes were directly obtained from Twist Bioscience already cloned into the pET28(a) expression vector. All expression plasmids were subsequently transformed into *E. coli* BL21(DE3) cells for protein expression.

### Protein expression

A secondary culture (500 mL LB broth (50 µg/mL kanamycin)) of *E coli*. BL21(DE3) encoding the protein of interest was expanded from an overnight primary culture (3 mL) in an incubator shaker at 37 °C until OD600 reached about 0.6 (⁓ 4hours). Protein expression was induced by addition of an IPTG solution to a final concentration of 1 mM. The induced culture was incubated in an incubator shaker for 2 h at 37 °C after which the culture was centrifuged and the bacteria pellet was frozen and stored in a -80 °C freezer. For purification, the bacteria pellet was thawed in a 37 °C water bath and lysed with a B-PER™ complete bacterial protein extraction reagent (ThermoFisher). The lysed solution was centrifuged and the supernatant was incubated with Qiagen Ni-NTA agarose for 30minutes at 4 °C to capture the product. The agarose was transferred to a fritted column, washed sequentially with a wash buffer A (50 mM Tris, 500 mM NaCl, 2% Triton X-114) (10 column volumes (CVs)) and a wash buffer B (50 mM Tris, 500 mM NaCl) (10 column volumes) before elution with an elution buffer (500 mM imidazole, 50 mM Tris, 500 mM NaCl, 10% glycerol) (3 CVs). The purified protein was confirmed via SDS-PAGE analysis with Coomassie blue staining.

### Synthesis of Conjugate Azide-AAPV-calicheamicin

An N-terminal azidoacetylated peptide substrate with a C-terminal penicillamine (Pen) (Azide-AAPV-Pen) was synthesized by CPC Scientific via a standard Fmoc-based solid phase peptide synthesis. A solution of calicheamicin (1 eq.) (MedChemExpress) in N,N-dimethylformamide (DMF) was added to a solution of Azide-AAPV-Pen (3 eq.) in DMF followed by addition of triethylamine (8 eq.) to the reaction solution. The reaction solution was purged with nitrogen for 5minutes and incubated for 24hours at 4 °C. The Azide-AAPV-calicheamicin product was purified by the Koch Institute Biopolymers & Proteomics core facility using a semi-preparative HPLC column and confirmed with MALDI-ToF MS. (ABD)_2_-(EEG)_6_-Cys was incubated with TCEP (10 eq.) for 30minutes at RT and buffer-exchanged into PBS (1 mM EDTA, pH 6.5) using an Amicon 10-kDa, ultra-0.5 centrifugal filter unit (14,000 rpm, 2minutes, 4 times). The reduced protein was incubated with DBCO-Maleimide (4 eq.) for 4hours at RT. The ABD-DBCO product was purified and buffer-exchanged into PBS using a PD-10 desalting column (Cytiva).

#### Synthesis of ABD-calicheamicin conjugates

Azide-Sx-calicheamicin (2 eq.) was dissolved in DMF and added to a solution of ABD-DBCO (1 eq.) in PBS to a final concentration of 50% DMF. The reaction solution was incubated for 24hours at RT and centrifuged to remove precipitate. The product in supernatant was immobilized on Ni-NTA agarose, washed with a wash buffer C (50 mM Tris, 500 mM NaCl, 20% DMF) (10 CVs) and a wash buffer B (10 CVs) before elution with the elution buffer. The product was buffer-exchanged into PBS using a PD-10 desalting column and quantified via absorbance at 200 nm. The product was confirmed and assessed for percentage purity using SDS-PAGE analysis.

### *In vitro* activity and toxicity masking

To evaluate conditional activation, ABD–calicheamicin conjugates were incubated with recombinant human neutrophil elastase (Enzo Life Sciences) or vehicle control in PBS at 37 °C for 4 h. Reaction mixtures were then diluted into bacterial or mammalian assay media as described below. Minimum inhibitory concentrations (MICs) were determined in accordance with CLSI guidelines. Briefly, bacteria were seeded at OD 0.005 in 96-well plates containing MHB supplemented with conjugate, protease-treated conjugate, or controls. Plates were incubated at 37 °C for 24 h, and bacterial growth was assessed by measuring optical density at 600 nm (SpectraMax i3, Molecular Devices). MICs were defined as the lowest concentration of conjugate preventing visible bacterial growth. Human hepatocellular carcinoma cells (HepG2, ATCC HB-8065) were maintained in Dulbecco’s Modified Eagle Medium (DMEM; Thermo Fisher Scientific) supplemented with 10% fetal bovine serum (FBS) and 1% penicillin– streptomycin. For cytotoxicity assays, HepG2 cells were seeded at 1 × 10^4 cells per well in collagen-coated 96-well plates and allowed to adhere overnight. Cells were then exposed to intact or protease-treated ABD–calicheamicin conjugates for 24 h. Viability was quantified using the CellTiter AQueous MTS assay (Promega) according to the manufacturer’s instructions, and absorbance was read at 490 nm using a plate reader. In vitro activity and toxicity data were fit using a four-parameter logistic (4PL) sigmoidal model, where X represents concentration, in GraphPad Prism (GraphPad Software).

### *In vitro* CalU19 neutralization evaluation

*SDS-PAGE* CalU19 was labeled with Cy7 through random lysine labeling and incubated with vehicle, calicheamicin, or ABD-Cal for 4 hours at 37C. At 4h, reactions were run on a gel (NuPAGE™ 4-12% Bis-Tris Protein Gels, 1.0 mm, 12-well, Thermofisher) which was then imaged using an Odyssey CLx imager in the 800 nm channel, for detection of Cy7 fluorescence, monitoring specifically for the presence of the intact protein at ∼27 kDa, as well as the self-cleavage fragments at 6kDa and 21 kDa.

*Cytotoxicity Assay* HepG2 cells were seeded at 1 × 10^4 cells per well in collagen-coated 96-well plates and allowed to adhere overnight. Cells were then exposed to serial dilutions of calicheamicin, in the presence of absence of 5 µM CalU19, CalU19x, C-CalU19X, LC-CalU19x, or vehicle (for the control). Viability was quantified at 24 h using the CellTiter AQueous MTS assay (Promega) according to the manufacturer’s instructions, and absorbance was read at 490 nm using a plate reader. Data were fit using a four-parameter logistic (4PL) sigmoidal model, where X represents concentration, in GraphPad Prism (GraphPad Software).

### *In vivo* pharmacokinetics, activity and toxicity evaluation

All animal studies were approved by the Massachusetts Institute of Technology’s Committee on Animal Care (MIT CAC protocol 2203000310). For infected mice, bacterial inoculum was prepared from mid-logarithmic phase cultures grown in Mueller–Hinton broth, washed twice with PBS, and quantified by optical density and serial dilution plating. Following administration, mice were monitored continuously until recovery from anesthesia and then observed at regular intervals for clinical signs of infection, including weight loss, respiratory distress, and reduced activity. For lung infections, CD-1 mice (10-12 weeks old) were administered 1.25 x 10^8^ CFU of *S. aureus* (502A) (suspended in PBS) or 5 x 10^6^ CFU of *P. aeruginosa* (suspended in PBS) by intratracheal administration using a 22G blunt end catheter and monitored after administration. At designated time points post-infection, mice were euthanized, lungs were aseptically harvested, and/or blood was collected.

*CFU quantification* Lungs were weighed and homogenized in sterile PBS using gentleMACS M-tubes (Miltenyi Biotec). Homogenates were serially diluted and plated on mannitol salt agar (BD Difco) to selectively recover *S. aureus*, and Pseudomonas Isolation Agar (Sigma-Aldrich) with colonies were enumerated after 24 h incubation at 37 °C. Samples with no detectable colonies on plates were recorded as below the limit of detection. For graphical representation and statistical comparison, these samples were assigned a value equal to the limit of detection.

*Histology* Lungs were fixed in 10% neutral buffered formalin for at least 48 h, processed, and paraffin-embedded by the Koch Institute Histology Core Facility. Tissue sections (X µm) were stained with hematoxylin and eosin (H&E) and imaged using brightfield microscopy. Histological evaluation was performed by

*Toxicity evaluation* To evaluate liver toxicity, blood was collected from mice at designated time points via terminal cardiac puncture. Whole blood was allowed to clot at room temperature in BD Microtainer SST tubes with gold tops (BD Biosciences, Cat. No. 365967). The resulting serum was aliquoted and stored at –80 °C until analysis. Serum alanine aminotransferase (ALT) activity was quantified using a colorimetric assay kit (Thermo Fisher Scientific, ALT/GPT Activity Assay Kit, Cat. No. EEA001, and serum aspartate aminotransferase (AST) activity was measured using a colorimetric assay kit (MilliporeSigma, AST Activity Assay Kit, Cat. No. MAK055), following the manufacturers’ protocols. Absorbance was read using a microplate reader (SpectraMax i3, Molecular Devices).

*Pharmacokinetics* CalU19 variants were conjugated with Cy7 through random lysine labeling. Labeled proteins were administered intravenously to CD-1 mice. At the indicated time points, blood was collected from the tail vein using heparinized capillary tubes, diluted in PBS containing EDTA, and centrifuged to remove cells. Fluorescence in plasma supernatants was quantified using a microplate reader to determine circulating protein concentrations. Percentage of antidote protein remaining in circulation was calculated as the percentage of antidote at each timepoint relative to the 1 min timepoint.

### Statistical Analysis and Schematic Representation

Statistical analyses were performed with GraphPad Prism. Data are plotted as mean ± SD. Comparisons among different treatment groups are based on one-way ANOVA with Tukey *post hoc* tests. A *P* value < 0.05 was considered statistically significant. Parts of the schematics in this publication were created with BioRender.

## SUPPLEMENTARY FIGURES

Figure S1. H&E-stained liver sections from mice receiving calicheamicin vs. ABD–Cal

Figure S2. CalU19 neutralizes free calicheamicin but not ABD-Cal

Figure S3. H&E-stained liver sections from mice receiving calicheamicin with or without LC-CalU19x.

## Notes

### Competing Interest Statement

S.N.B. reports consulting roles and/or equity Amplifyer Bio, Catalio Capital, Danaher, Earli Inc., Impilo Therapeutics, Matrisome Bio, Ochre Bio, Port Therapeutics, Satellite Bio, Sunbird Bio, and Xilio Therapeutics, which were not involved in this study.

### Summary of Updates

Abstract, Introduction, Results and Discussion updated, additional data included.

