## Supporting Information for "Expanding antimicrobial chemical space by engineering drug safety"

Supplementary Figure 1

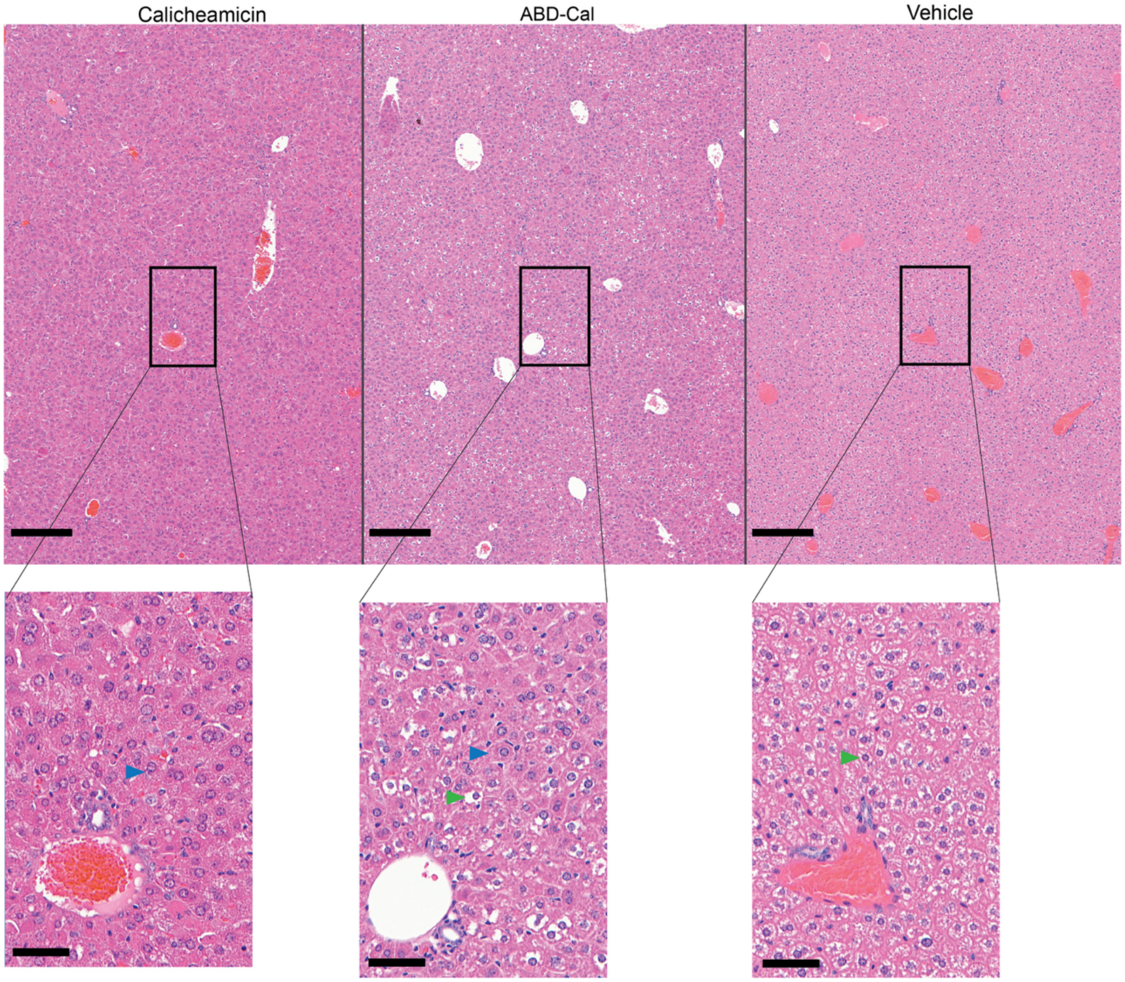


**Supplementary Figure 1 |** Representative hematoxylin and eosin (H&E) stained liver sections from mice receiving calicheamicin (0.025 mg/kg), ABD–Cal (0.025 mg/kg drug eq.), or vehicle (control). Scale bar = 200 μm, scale bar of inset = 50 μm. Blue arrows indicate swollen hepatocytes with eosinophilic cytoplasm, green arrows indicate healthy hepatocytes.

Supplementary Figure 2

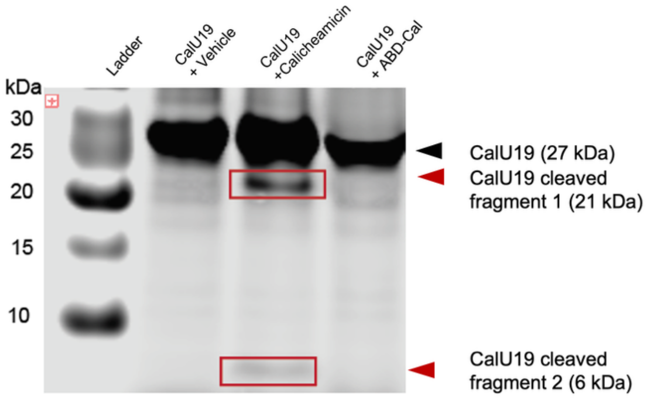


**Supplementary Figure 2 | CalU19 neutralizes free calicheamicin but not ABD-Cal.** Assessment of **n**eutralization of free calicheamicin versus conjugated calicheamicin (ABD-Cal) by CalU19, assessed by SDS–PAGE of Cy7-labeled CalU19. CalC, CalU19, and CalU16 are known to undergo a unique self-cleavage event in the process of calicheamicin neutralization**.** CalU19 incubated with vehicle shows intact CalU19 (27 kDa); CalU19 incubated with free calicheamicin shows CalU19 self-cleavage fragments (21 kDa and 6 kDa); CalU19 incubated with ABD-Cal shows intact CalU19, but not self-cleavage fragments.

Supplementary Figure 3

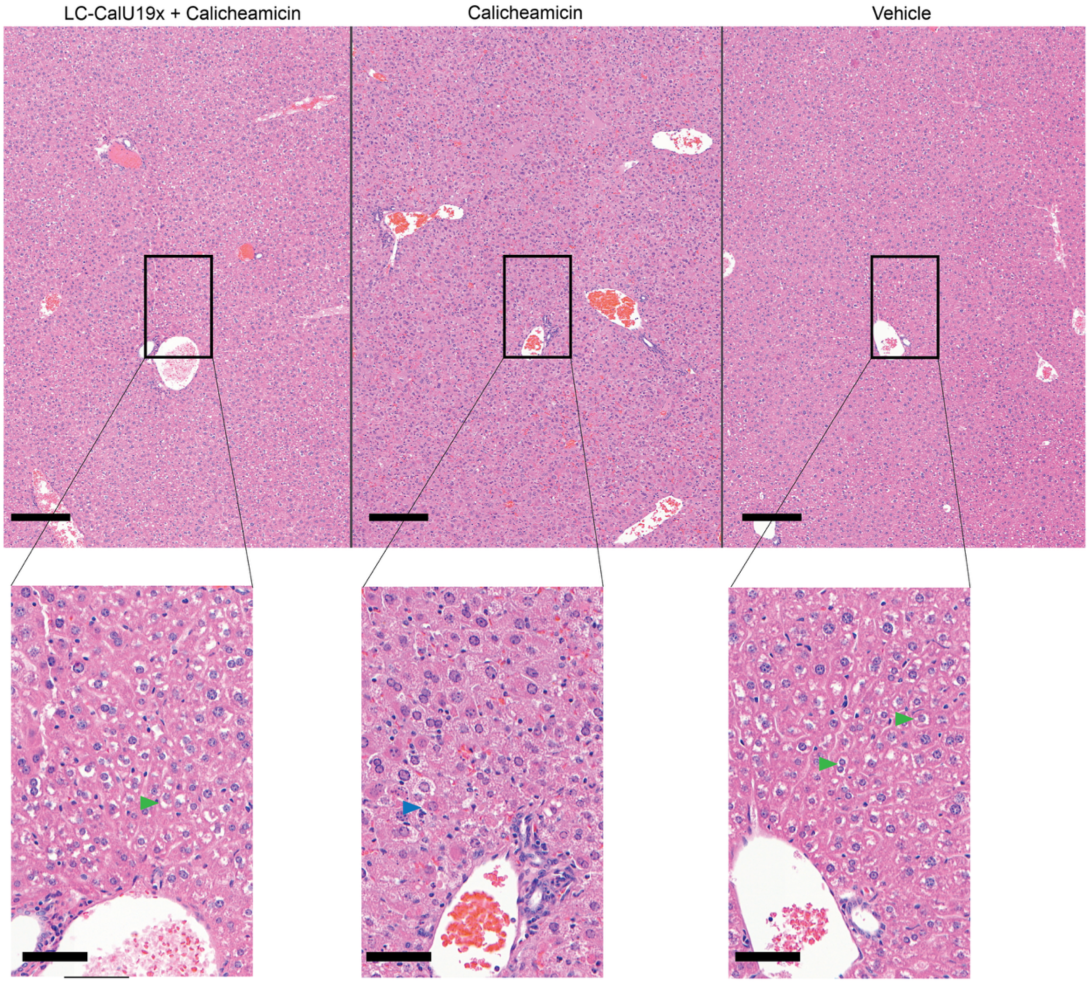


**Supplementary Figure 3 | R**epresentative hematoxylin and eosin (H&E) stained liver sections from mice receiving calicheamicin (0.025 mg/kg) with or without LC-CalU19X. Representative image of mice receiving vehicle included as control. Scale bar = 200 μm, scale bar of inset = 50 μm. Blue arrows indicate swollen hepatocytes with eosinophilic cytoplasm, green arrows indicate healthy hepatocytes.
